# Chilling induces unidirectional solute leak through the locust gut epithelia

**DOI:** 10.1101/784504

**Authors:** Kaylen Brzezinski, Heath A. MacMillan

## Abstract

Chill-susceptible insects, like the migratory locust, often die when exposed to low temperatures from an accumulation of tissue damage that is unrelated to freezing (chilling injuries). Chilling injury is consistently associated with ion imbalance across the gut epithelia. It has recently been suggested that this imbalance is at least partly caused by a cold-induced disruption of epithelial barrier function. Here, we aim to test this hypothesis in the migratory locust (*L. migratoria*). First, chill tolerance was quantified by exposing locusts to −2°C for various durations and monitored for chill coma recovery time and survival 24h post-cold exposure. Longer exposure times significantly increased recovery time and caused injury and death. Ion-selective microelectrodes were also used to determine the presence of cold-induced ion imbalance. We found a significant increase and decrease of hemolymph K^+^ and Na^+^ concentrations over time, respectively. Next, barrier failure along the gut was tested by monitoring the movement of an epithelial barrier marker (FITC-dextran) across the gut epithelia during exposure to −2°C. We found minimal marker movement across the epithelia in the serosal to mucosal direction, suggesting that locust gut barrier function remains generally conserved during chilling. However, when tested in the mucosal to serosal direction, we saw significant increases of FITC-dextran with chilling. This instead suggests that while cold-induced barrier disruption is present, it is likely unidirectional. It is important to note that these data reveal only the phenomenon itself. The location of this leak as well as the underlying mechanisms remain unclear and require further investigation.

**Summary statement:** In this study, we provide the first evidence for the presence of cold-induced paracellular leak along the gut of the migratory locust, and that this leak is strongest in the mucosal to serosal direction.

## Introduction

Chill susceptible insects are those that succumb to the negative effects of cooling at temperatures well above the freezing point of their extracellular fluids (Overgaard and MacMillan, 2017). When many ectothermic animals are chilled, they reach a temperature at which a state of neuromuscular paralysis (chill coma) occurs, known as chill coma onset temperature (CCO). Insects can remain in this reversible comatose state for the duration of a cold exposure in the lab and can often recover once the stressor (brief or mild cold) has been removed. The time taken to regain the ability to stand following chill coma is termed chill coma recovery time (CCRT) (Gibert et al., 2001; David et al., 1998). Both CCO and CCRT are regularly used as non-lethal means of quantifying chill susceptibility in various groups of insects, such as crickets (*Gryllus pennsylvanicus*), caterpillars (*Pringleophaga marioni*), fruit flies (*Drosophila melanogaster*), locusts (*Locusta migratoria*), and firebugs (*Pyrrhocoris apterus*) (Andersen et al., 2013; Andersen et al., 2017a; Chown and Klok, 1997; Coello Alvarado et al., 2015; Robertson et al., 2017). In the event of a particularly harsh or prolonged exposure, an accumulation of cold-induced tissue damage (chilling injuries) can occur (Coello Alvarado et al., 2015; Koštál et al., 2006; MacMillan and Sinclair, 2011a). Chilling injuries typically manifest as a loss of coordination, permanent limb paralysis, or mortality (Overgaard and MacMillan, 2017). Quantifying an insect’s condition or survival following cold stress is therefore another common measure of chill tolerance, and can be quantified using a scoring system (e.g. as dead/alive, or a range of conditions from dead to alive) to indirectly measure the degree of injury sustained (Findsen et al., 2013; Overgaard and MacMillan, 2017). While each of these chill tolerance traits operate via distinct mechanisms, they have all been in some manner associated with a local or systemic loss of ion balance, suggesting that ionoregulatory failure may be a principal cause of chill susceptibility (Armstrong et al., 2012; Bayley et al., 2018; Findsen et al., 2013; Koštál et al., 2006; MacMillan et al., 2015a; Robertson et al., 2017).

In insects, organismal ion homeostasis is under tight regulation by both the Malpighian tubules (MTs) and gut epithelia (MacMillan and Sinclair, 2011a). Briefly, the gut has three main regions (foregut, midgut, and hindgut), each with its own specialized functions. The foregut lies at the anterior margin of the gut and consists of flattened and undifferentiated cells consistent with the lack of absorption or secretion that takes place in the region (Chapman, 2013). Instead, the foregut generally acts as a passage through which food travels, although muscular breakdown and salivary digestion of the bolus can occur (Dadd, 1970). From the foregut, food travels to reach the remainder of the gut where it is digested (Engel and Moran, 2013; Naikkhwah and O’Donnell, 2012). Cells along the midgut are actively involved in digestive enzyme production and secretion, acting as the primary site of digestion and nutrient, ion, and water absorption (Chapman, 2013; Yerushalmi et al., 2018). Following the midgut are the MTs, blind-ended tube-shaped diverticula of the gut that are somewhat analogous to the vertebrate kidneys (Chapman, 2013). These tubules are one cell layer thick and contain a multitude of cation exchangers and pumps (Maddrell and O’Donnell, 1992; O’Donnell and Ruiz-Sanchez, 2015). Through these ionoregulatory pumps (e.g. V-ATPase, the proton pump that primarily energizes transport at the MTs) and channels, ions such as K^+^, Na^+^ and Cl^-^ are driven from the hemolymph to the tubule lumen. For instance, K^+^ is secreted into the tubule lumen which helps to maintain a low K^+^ (generally 10-15 mM) environment that permits muscle function (Andersen et al., 2017a; Andersen et al., 2018; Djamgoz, 1987; Gerber and Overgaard, 2018; Harvey and Zerahn, 1972; Hoyle, 1953; Rheuben, 1972; Yerushalmi et al., 2018). These means of ionoregulation across the MTs promote an osmotic gradient that favours the movement of water and unwanted waste products or toxins into the tubule lumen (secretion), producing primary urine (Maddrell and O’Donnell, 1992). Finally, the hindgut (composed of the ileum and rectum) is the most posterior region of the gut, and is the main site for the absorption of water and solutes (Phillips et al., 1987). Cell membranes along the intercellular spaces of the hindgut are rich in Na^+^/K^+^-ATPase, which generates high [Na^+^] in the paracellular space that drives water across the hindgut epithelium via osmosis, permitting water reabsorption and production of a dry feces (Des Marteaux et al., 2018; Phillips, 1981; Wall and Oschman, 1970).

Through the continuous ionoregulatory actions of the alimentary canal, the hemolymph of most insects contains high and low concentrations of Na^+^ and K^+^, respectively, at optimal or near-optimal thermal conditions (Engel and Moran, 2013; MacMillan et al., 2015b; Maddrell and O’Donnell, 1992). In cold conditions, however, the activity of ionoregulatory enzymes like V-ATPases and Na^+^/K^+^-ATPases is suppressed (Bayley et al., 2018; Hosler et al., 2000; Mandel et al., 1980; McMullen and Storey, 2008; Moriyama and Nelson, 1989; Yerushalmi et al., 2018). Over time, this temperature-induced suppression of transcellular ion transport often results in a net leak of hemolymph Na^+^ and water (which follows Na^+^ osmotically) to the gut lumen, effectively concentrating the K^+^ that remains in the hemolymph (MacMillan and Sinclair, 2011b). Some intracellular K^+^ simultaneously leaking down its concentration gradient into the extracellular space worsens this problem (Andersen et al., 2017; MacMillan et al., 2014). As K^+^ concentrations rise in the hemocoel (hyperkalemia), the gradient of K^+^ across the cell membrane is lost, and a marked depolarization in membrane potential occurs, eventually resulting in the activation of voltage-gated Ca^2+^ channels (Andersen et al., 2017a; Bayley et al., 2018; MacMillan et al., 2015a). The influx of Ca^2+^ that ensues is proposed to initiate a crippling cascade which causes a deterioration of cellular integrity, likely through apoptosis (Mattson and Chan, 2003; Nicotera and Orrenius, 1998; Yi and Lee, 2011; Yi et al., 2007). Failure to maintain ion and water homeostasis in the cold can therefore ultimately result in organismal chilling injury or death. In turn, understanding the biochemical mechanisms underlying this failure is critical to understanding chill susceptibility. While a cold-induced failure of transcellular transport is one likely mechanism of chilling injury and is under active investigation, ions do not cross epithelia solely via transcellular pathways (Donohoe et al., 2000; MacMillan et al., 2016a; O’Donnell and Maddrell, 1983).

In addition to the active movement of ions and passive transport of solutes and water through cells (transcellular pathways; e.g. via channels), the gut also relies on the passive movement, or leak, of molecules through paracellular pathways between adjacent cells (Jonusaite et al., 2016; le Skaer et al., 1987). Septate junctions (SJs) are specialized cell-cell junctions analogous to vertebrate tight junctions that largely determine the permeability of these paracellular pathways (Jonusaite et al., 2016). Arthropod epithelia generally have two types of SJs: pleated and smooth. The former are typically observed in ectodermally-derived epithelia such as the foregut and hindgut, and are 2-3 nm wide, while smooth SJs are found in endodermally-derived tissues like the midgut, and are 5-20 nm wide (Jonusaite et al., 2016). To date, the majority of SJ studies has been conducted on *Drosophila*, including the identification of SJ types, associated proteins, and SJ influence in cold tolerance plasticity (Izumi and Furuse, 2014; Jonusaite et al., 2016; MacMillan et al., 2017). Notably, flies acclimated to colder conditions are more cold tolerant than warm-acclimated flies and upregulate approximately 60% of genes encoding known or putative fly SJ proteins (MacMillan et al., 2016b). These cold-acclimated (10°C) flies also have reduced rates of paracellular leak of a fluorescent probe (FITC-dextran) from their gut lumen to their hemolymph, both before and during a cold stress, compared to warm-acclimated (25°C) flies (MacMillan et al., 2017). Together, these studies suggest that cold exposure can cause increased rates of leak through the paracellular barriers, and that improvements in cold tolerance may be in-part related to an improved ability to maintain paracellular barriers. However, because flies were fed the probe for these experiments (and the gut was completely loaded with the probe upon cold exposure) the precise location of this leak along the gut and the mechanisms that drive it remain unclear, as does whether this a problem experienced by all insects, or just *D. melanogaster*.

Here, we investigate the effects of chilling on epithelial barrier integrity in a chill susceptible insect, the migratory locust (*Locusta migratoria*). As previously observed in *Drosophila*, we hypothesized that chilling disrupts septate junctions (SJs) in the locust gut and that this effect leads to paracellular leak across the gut epithelia. Due to the often temperature-sensitive nature of ionoregulatory enzymes, and their dense concentration along ionomotive epithelia, we also hypothesized that this disruption in barrier integrity is limited to transport-rich segments along the locust gut such as the midgut and hindgut. To address these hypotheses, we first confirmed that our locust colony is chill susceptible by measuring their survival and performance post-cold exposure. We then used the fluorescently-labelled marker, FITC-dextran to characterize directionality in cold-induced paracellular leak, and find that cold does induce paracellular leak through the gut epithelia of locusts, but surprisingly only in one direction.

## Methods

### Experimental system

All experiments were conducted using male and female adult locusts (*Locusta migratoria*) aged 3-4 weeks post-final ecdysis. Locusts were obtained from a continuously breeding colony maintained at Carleton University in Ottawa, ON. This colony is reared under crowded conditions on a 16 h:8 h light:dark cycle at 30°C with 60% relative humidity (see Dawson et al., 2004). All animals had *ad libitum* access to a dry food mixture (oats, wheat bran, wheat germ, and powdered milk), and fresh wheat clippings supplied three times per week.

### Chill coma recovery time and survival

The degree of chill susceptibility of the locust colony was quantified using both their chill coma recovery time (CCRT) and the degree of injury/mortality 24 h following exposure to −2°C. On the day of the experiment, locusts were collected from the colony, sexed by eye, and individually placed into 50 mL polypropylene tubes. These tubes were sealed using lids with small holes to allow access to air for the duration of the experiment. Excluding controls, all locusts were suspended using a Styrofoam rig in a cooling bath (Model AP28R-30, VWR International, Radnor, USA) containing a circulating ethylene glycol:water mix (3:2) preset to 20°C and cooled to −2°C at a rate of −0.20°C min^−1^. Both bath temperature and locust internal body temperature were monitored; the former via internal probes, and the later via inserted type-K thermocouples (TC-08 interface; *PicoLog* software version 5.25.3) located at the junction of the head and thorax of representative locusts (that were not used further in the experiments). Locusts were then left undisturbed for 2, 6, 24, or 48 h upon which arbitrarily selected groups of locusts were removed from the cold and returned to room temperature (23°C). In order to monitor CCRT, insects were removed from their tubes and gently placed on the surface of a table and observed for the time taken to recover from chill coma. Animals were stimulated by gentle puffs of air from a transfer pipette every minute and were marked as having fully recovered when standing on all six limbs. Observation time was limited to 60 min; any locusts not meeting this criterion were marked as having not recovered.

After 60 min, the locusts were returned to their respective tubes along with a dry food mixture (oats, wheat bran, wheat germ, and powdered milk) and water (supplied in microcentrifuge tubes with cotton) and left for 24 h at room temperature (23°C). An assessment of 24 h survival post-cold exposure was performed using a scoring system of 0 to 5, similar to that described by MacMillan et al. (2014). Briefly, scores were defined as follows: 0: no movement observed (i.e. dead); 1: limb movement (slight leg and or head twitching); 2: greater limb movement, but unable to stand; 3: able to stand, but unable or unwilling to walk or jump; 4: able to stand, walk, and or jump, but lacks coordination; and 5: movement restored pre-exposure levels of coordination.

### Quantification of serosal to mucosal paracellular leak and ion imbalance

To measure paracellular permeability in the gut epithelia of locusts, we monitored the movement of a fluorescently-labeled molecule in both the serosal (hemolymph) to mucosal (lumen) direction, and the mucosal to serosal direction. All experiments used FITC-dextran (3-5kDa, Sigma Aldrich, St. Louis, USA) a commonly used probe for determining paracellular permeability in both invertebrate and vertebrate models such as fruit flies (*D. melanogaster*), rats (*Rattus norvegicus domesticus*), and zebra fish (*Danio rerio*) (see MacMillan et al., 2017; Condette et al., 2014; Bagnat et al., 2007).

Experiments conducted in the serosal to mucosal direction (from the hemolymph to the gut lumen) were done both to identify the presence of leak across the gut epithelia and isolate the area across which leak occurred. Protocols for this novel leak assay were developed through preliminary trials. In the final assay, FITC-dextran was dissolved in locust saline (in mmol 1^−1^: 140 NaCl, 8 KCl, 2.3 CaCl_2_ Dihydrate, 0.93 MgCl_2_ Hexahydrate, 1 NaH_2_PO_4_, 90 sucrose, 5 glucose, 5 trehalose, 1 proline, 10 HEPES, pH 7.2) resulting in a final FITC-dextran concentration of 3.84 x 10^-3^ M (selected based on standard curves from the preliminary trials). Using a 25 μL Hamilton syringe, 20 μL of this solution was injected into the hemocoel ventrally at the junction of the thorax and first abdominal segment of locusts. Pilot experiments revealed that neither the 20 μL injection nor the FITC-dextran itself impacted locust performance or survival (survival for control locusts with and without FITC-dextran injection was scored as 5). Following the protocol for CCRT (outlined above), animals were suspended in a cooling bath preset to 20°C and cooled to −2°C at a rate of −0.2°C min^−1^. Insects were then held at −2°C (or room temperature for controls) and left undisturbed for 2, 6, 24, or 48 h.

Locusts were individually removed from the cooling bath and dissected within 15 min of their target cold exposure duration for tissue collection. Animals were sacrificed by decapitation before removing all limbs and wings. The thorax and abdomen (containing the internal organs) were placed in a petri dish lined with silicone elastomer (Sylgaard 184 Silicone Elastomer Kit, Dow Chemical, Midland, USA) and containing locust saline. A longitudinal incision was made in the anterior to posterior direction along the ventral side to expose the gut. With the body wall pinned back, the tracheae and Malpighian tubules were then cleared away to access the gut tissue. The gut was then cut into three segments (anterior, central, posterior) based on our ability to carefully isolate these segments rather than pre-determined anatomical divisions (see Fig. 2A). Briefly, the anterior segment was defined as the foregut to the anterior midgut caeca, the central segment as the posterior midgut caeca to the midgut-hindgut junction, and the posterior segment as the midgut-hindgut junction to the rectum. To avoid excessive leak of gut contents during collection, segments were gently pinched with dissecting forceps at both ends before excision. Upon removal, segments were washed briefly in saline (while retaining their contents) to remove any excess dextran-saturated hemolymph, and placed in microcentrifuge tubes containing 500 μL of locust saline. Samples were subsequently homogenized (OMNI International Tissue Master 125 120 V, Kennesaw, USA; approximately 3 min), sonicated (Qsonica Sonicators Model CL-188, Newton, USA; 3 x 5 s bursts with 10 s rests), and centrifuged for 5 min at 10,000 × *g*. A 100 μL aliquot of the resulting supernatant was collected and transferred to a black 96-well plate for fluorescence spectrophotometry (Ex: 485 nm, Em: 528 nm; BioTek Cytation 5 Imaging Reader, Winooski, USA). Concentrations of FITC-dextran in the samples were determined by reference to a standard curve of FITC-dextran in locust saline, and control samples confirmed that tissues from locusts that were not injected with the probe had negligible fluorescence (see Fig. 2B and D).

Because little FITC-dextran appeared in the gut samples (see Results), hemolymph extraction experiments were performed on separate locusts over identical exposures to determine how much FITC-dextran was being lost from the hemolymph over time. Similar to the above protocols, a new set of locusts were injected with the FITC-dextran solution and suspended at - 2°C for 2, 6, 24, or 48 h in a circulating cooling bath, while controls were held at room temperature. After their designated exposures, hemolymph samples were collected using methods adapted from Findsen et al. (2013). Briefly, locusts were pricked dorsally using a dissecting probe at the junction of the head and thorax before using a 50 μL capillary tube to collect the hemolymph (as described for hemolymph ion measurements). A 2 μL aliquot of hemolymph was pipetted into 96-well plates (Corning Falcon Imaging Microplate; black/clear bottom), diluted 50-fold with saline, and analyzed for FITC-dextran content via fluorescence spectrophotometry. Pilot experiments showed no interference from the saline when measuring fluorescence. To maximize use of the large volumes of available hemolymph, these locusts were also used to collect data on hemolymph ion balance over time.

An additional 10 μL of hemolymph from each animal was collected using a 50 μL capillary tube. Samples were promptly vortexed and flash frozen in liquid nitrogen to avoid coagulation of the hemolymph and stored at −80°C until experiments. All samples were vortexed once again prior to testing. Hemolymph Na^+^ and K^+^ concentrations were measured using ion-selective borosilicate microelectrodes (TWI150-4, World Precision Instruments, Sarasota, USA). No interference from the FITC-dextran was found when measuring ions. The ion content of hemolymph both with and without FITC-dextran was measured in control (room temperature) locusts across 48h, and statistical analysis revealed no significant differences in ion concentrations between the two groups (Linear model, *F_1, 10_* = 0.302, *P* = 0.594). Our Na^+^-selective microelectrodes were constructed using 100 mM NaCl backfill solution and Na^+^ ionophore II cocktail A (Sigma Aldrich), while K^+^ selective microelectrodes contained 100 mM KCl backfill solution and K^+^ ionophore I cocktail B (Sigma Aldrich). Microelectrodes were calibrated using standards of 10 mM and 100 mM of NaCl or KCl (osmolality adjusted with LiCl) for their respective measurement. These standards were also used to calculate the ion concentration in samples of hemolymph from the obtained voltage measurements using the following equation:

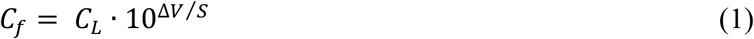

where *C_f_* is the final concentration in mM, *C_L_* is the concentration (in mM) of the lowest standard used for the data point of interest, Δ*V* is the difference (mV) between the sample of interest and the lowest standard, and *S* is the slope (the difference in mV between the two standards). Only microelectrodes with a slope between 50 and 60 mV (close to the expected Nernst slope of 58 mV) were used for all experiments (Na^+^: 51.2 ± 0.1; K^+^ 55.5 ± 0.4).

### Measuring paracellular leak in fed locusts

Hemolymph extractions were also used (on a separate set of locusts) to measure leak across the gut epithelia in the mucosal to serosal direction. Looking back on the literature, we noticed that experiments using models such as mosquitos (*Aedes aegypti* and *Anopheles gambiae*) and rats (*Rattus norvegicus domestica*) administered FITC-dextran orally to test for paracellular leak (Condette et al., 2014; Edwards and Jacobs-Lorena, 2000; Pantzar et al., 1993). To test whether the lack of FITC-dextran movement in the serosal to mucosal direction was due to this key difference in methodology, we took a different approach. Instead of FITC-dextran injections, we fed locusts a mixture of dry food (oats, wheat bran, wheat germ, and powdered milk) saturated with a solution of FITC-dextran in water (9.6 x 10^−4^ M) for 24 h prior to experiments. Pilot experiments confirmed the presence of FITC-dextran throughout the alimentary canal the following day. Similar to experiments in the opposite direction, all animals were exposed to −2°C for 2, 6, 24, or 48 h. Hemolymph was sampled and analyzed as above following removal from the cold.

### 2.2.6 Data Analysis

All data, excluding concentrations of FITC-dextran found in the gut (*in vivo* FITC-dextran injection experiments; outlined below), were analyzed using linear models (i.e. one- or two-way ANOVAs) in R Studio version 1.2.1335 (https://www.rstudio.com). The effects of time in the cold on gut leakiness (quantified by FITC-dextran movement) were analyzed using a linear mixed effects model via the lmer() function in R (lme4 and lmerTest packages for R). Time and segment were treated as fixed effects, while locust sex (*in vivo* FITC-dextran injection experiments) was treated as a random effect to account for variability in locust gut leakiness per individual or sex. All data were analyzed with time as both a continuous and categorical factor, however, the outcomes of these two approaches were identical. As such, all results presented in this section treat time as a continuous factor. The level of statistical significance was 0.05 for all analyses, while all additional values presented are mean ± standard error.

## Results

### Chill coma recovery time and injury following chilling

The chill susceptibility of our locust colony was confirmed by measuring chill coma recovery time and scoring injury/survival of locusts 24 h post-cold exposure. After 2 h of cold exposure, all animals recovered from chill coma to standing position (CCRT) within 10-18 min. However, recovery time significantly increased as exposure times grew longer (Fig. 1A; Linear model, *F_1, 22_* = 31.3, *P* < 0.001). This trend persisted until the last time point (48 h at −2°C), at which point no locusts recovered the ability to stand within 60 min. Similarly, survival rates decreased with longer cold exposures, leading to nearly 100% mortality after 48 h at −2°C (Fig. 1B; Linear model, *F_1, 38_* = 199, *P* < 0.001).

**Figure 1.**
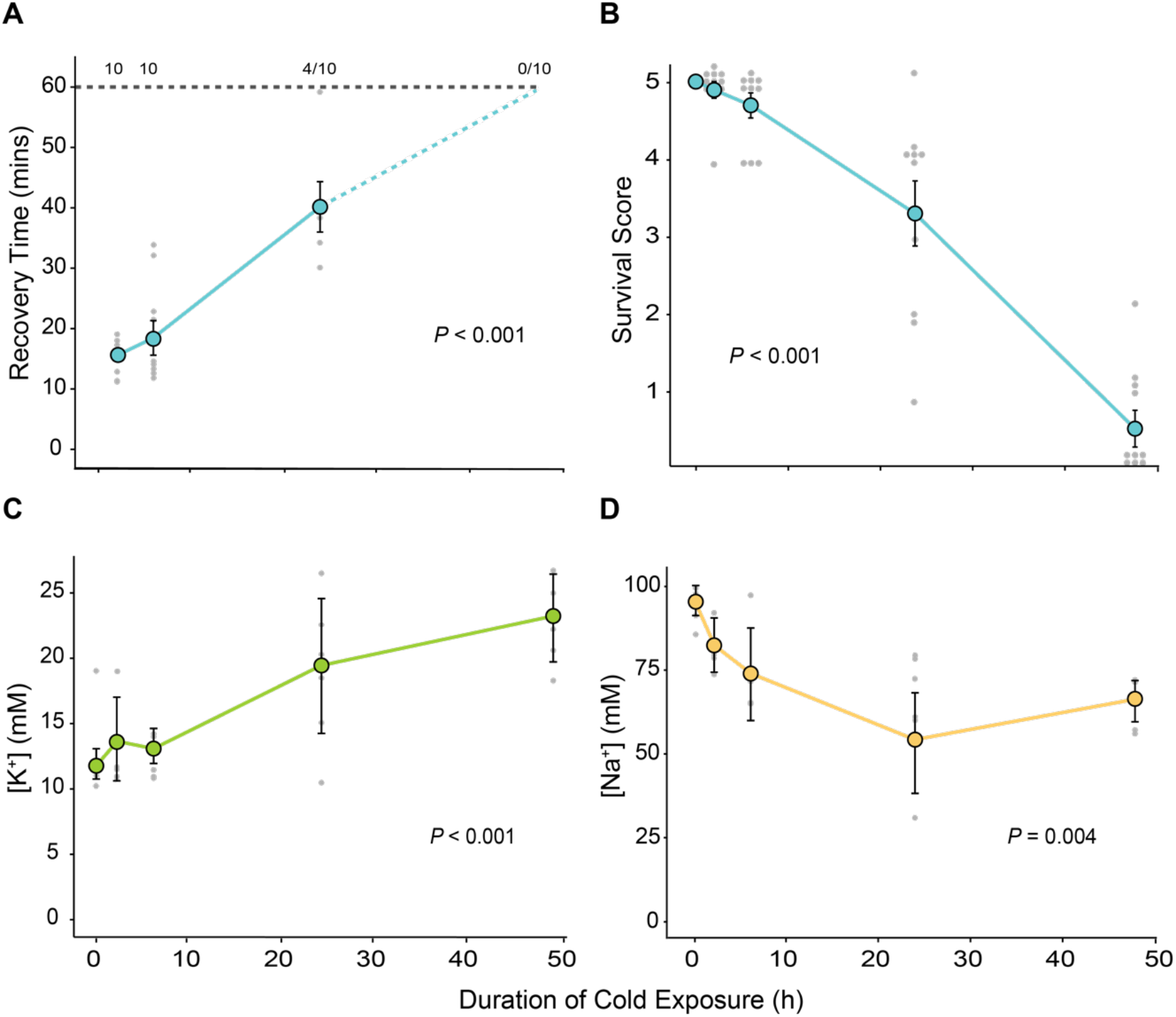
Locusts (*L. migratoria*) suffer from injury and ionoregulatory collapse typical of chill susceptible insects. **A)** Chill coma recovery time (CCRT) of locusts (mixed sexes; n = 10 per time point) held at −2 °C for 2, 6, 24, or 48 h. Locusts were observed for 60 mins following cold exposure and were marked as having recovered when standing on all six limbs. Values above the dotted black line represent the number of locusts that had recovered within 60 mins. The solid blue line represents mean values per time point. **B)** Locust condition (survival) following exposure to −2°C for 0, 2, 6, 24, or 48 h (n = 10 per time point). Survival score was based on the following: 0: no movement observed (i.e. dead); 1: limb movement (leg and or head twitching); 2: moving, but unable to stand; 3: able to stand, but unable or unwilling to walk or jump; 4: able to stand, walk, and or jump, but lacks coordination; and 5: movement restored pre-exposure levels of coordination. The solid blue line represents mean values per time point. To show all data points, dots are clustered around their respective score (where applicable). **C)** Changes in locust hemolymph K^+^ concentrations over time spent at −2°C (n = 5-6 locusts per time point). **D)** Samples of locust hemolymph Na^+^ concentrations over time spent at −2°C (n = 4-6 locusts per time point). Values are mean ± standard error. Light grey points represent each sample taken per time point. Error bars not shown are obscured by the symbols.

### Hemolymph ion concentrations

Ion selective microelectrodes were used to determine the effects of cold exposure on extracellular ion balance over the course of our experiments. While the concentration of Na^+^ in the hemolymph decreased significantly as time at −2°C increased up until approximately 24 h (Fig. 1D; Linear model, *F_1, 23_* = 10.1, *P* = 0.004), hemolymph K^+^ concentrations significantly increased, doubling from 11.8 mM to 23.3 mM over 48 h at −2°C (Fig. 1C; Linear model, *F_1, 23_* = 27.8, *P* < 0.001).

### Serosal to mucosal leak

To quantify the presence of paracellular leak across the gut epithelia in the cold (from the hemocoel into the gut), samples were taken from each gut segment (Fig. 2A, anterior, central, and posterior) and analyzed for fluorescent content following marker injection. Interestingly, while FITC-dextran concentrations significantly increased in the gut over time (Fig. 2B; Linear model, *F_1, 77_* = 7.85, *P* = 0.006), less than 1% of the total injected marker appeared within the gut after 48 h in the cold. Using summary data from both the gut leak assay (0.019 ± 0.003 mg/mL) and hemolymph extraction experiments (following section; 2.05 ± 0.279 mg/mL), we estimate that approximately 0.93% of total FITC-dextran injected into the hemolymph leaked across the gut epithelia into the lumen during the entire 48 h cold exposure. This method also made it possible for us to isolate potential sites of barrier loss along the gut. However, in addition to the minute movement of FITC-dextran over time, there were no significant differences in marker concentration among the three gut segments (Linear model, *F_2, 77_* = 1.76, *P* = 0.179). Similarly, no significant interaction was found between the time spent at −2°C and the segment type (Linear model, *F_2, 77_* = 1.57, *P* = 0.214). Finally, total concentrations of FITC-dextran found in the gut did not significantly differ between treatment and control locusts at 24 h (Fig. 2C; Linear model, *F_4, 1_* = 0.531, *P* = 0.758). There was, however, a significant difference between the total concentration of FITC-dextran sampled at 0 h and 24 h in both treatments (Linear model, *F_1, 11_* = 25.4, *P* < 0.001).

**Figure 2.**
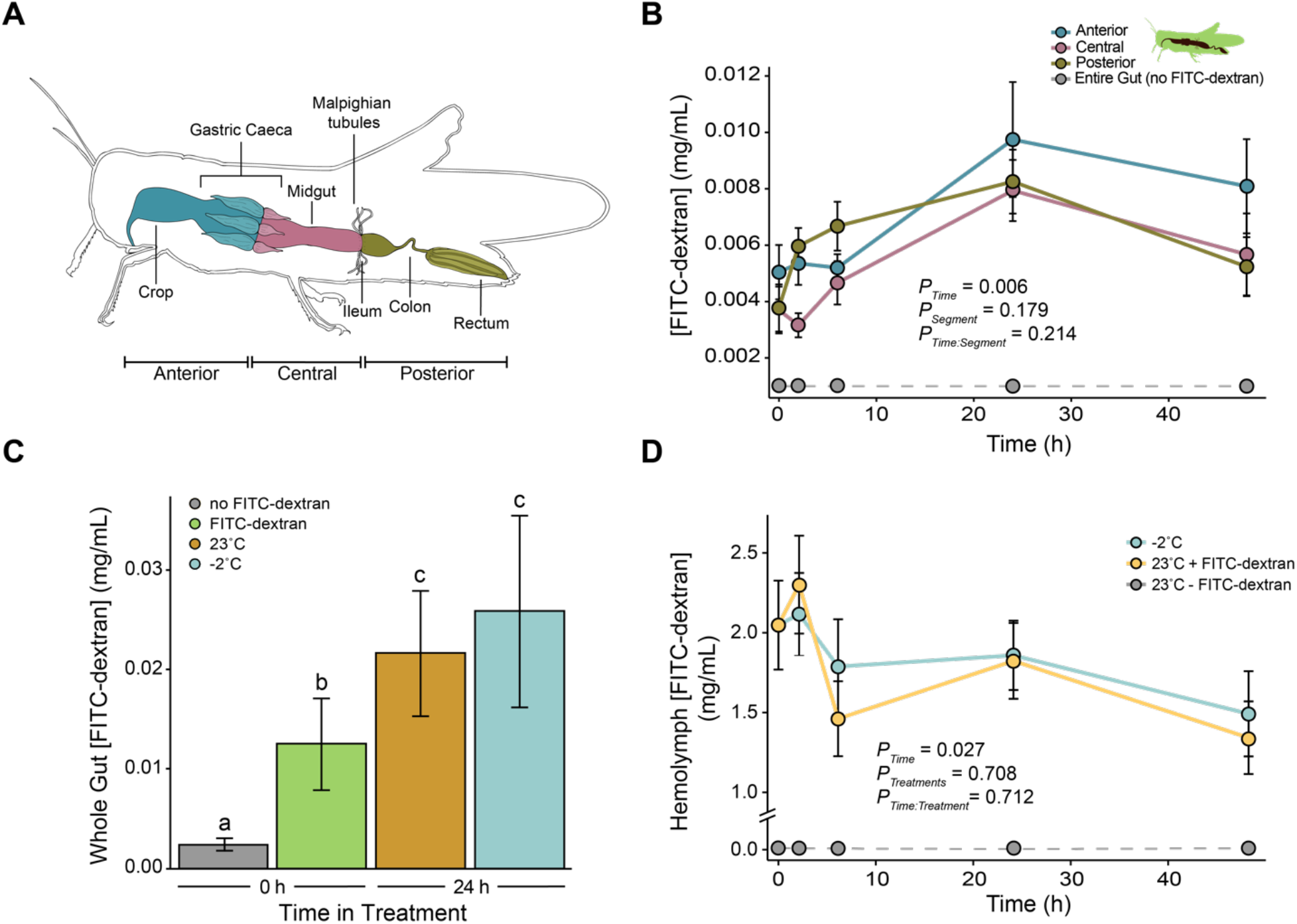
Cold stress causes minimal leak of FITC-dextran across the gut epithelia of *L. migratoria* in the serosal to mucosal direction. **A)** A schematic of the locust (*L. migratoria*) gut tract sectioned into three segments (anterior, central, and posterior). Relative to the locust gut anatomy, the segments were determined as follows: anterior – foregut to the anterior midgut caeca; central – posterior midgut caeca to the Malpighian tubules (removed; the midgut-hindgut junction); posterior – Malpighian tubules to the rectum. Figure illustrated from observation. **B)** Concentration of injected FITC-dextran (mg/mL) present in to each gut segment (anterior, central, and posterior; n = 4-6 per group) after exposure to −2°C for 2, 6, 24, or 48 h. **C)** Mean values representing the total FITC-dextran content within the gut (sum of anterior, central, and posterior segments) at 24 h of either 23°C (control) or −2°C (n = 4-6 per group). Means shown at 0 h were sampled immediately post-injection (FITC-dextran or saline). Letters denote a significant difference (*P_Time_* < 0.001; *P_Treatment_* = 0.758). **D)** Concentration of injected FITC-dextran remaining in the hemolymph over 48 h in the cold (−2°C) and at room temperature (n = 4-6 per group). Values are mean ± standard error. Error bars not shown are obscured by the symbols.

Loss of marker from the hemolymph was investigated to corroborate levels of cold-induced leak into the gut and to determine whether our locusts were capable of metabolizing the probe. There was a significant loss of FITC-dextran from the hemolymph over time, both in the cold and at room temperature (Fig. 2D; Linear model, *F*_1, 39_ = 5.26, *P* = 0.027). However, there was no significant difference in marker movement between the two treatment groups (Linear model, *F*_1, 39_ = 0.142, *P* = 0.708). We also observed no significant interaction between marker concentration over time in the cold and room temperature conditions (Linear model, *F_1, 39_* = 0.138, *P* = 0.712).

### Mucosal to serosal leak

Traditionally, studies examining paracellular leak of FITC-dextran and other large markers like inulin involve the oral administration of probes to the animals, which is in stark contrast to our serosal to mucosal (probe injection) approach. These differences in methods may account for our initial finding that paracellular barriers are maintained in the cold (at least in locusts). To address this possibility, we again examined cold-induced leak, however, this time in the mucosal to serosal direction. Locusts were fed a dry food mixture saturated with water containing a fixed concentration of FITC-dextran and sampled for marker content. Unlike the minimal FITC-dextran leak that occurred in the serosal to mucosal direction, these experiments yielded a significant and near linear increase in hemolymph FITC-dextran concentration over time in the cold (Fig. 3; Linear model, *F_1, 44_* = 10.8, *P* = 0.002). Finally, significant differences were also observed between treatment groups (Linear model, *F_1, 44_* = 9.40, *P* = 0.004). There was approximately a 15-fold difference in hemolymph FITC-dextran levels between control locusts and those that spent 48 h at −2°C.

**Figure 3.**
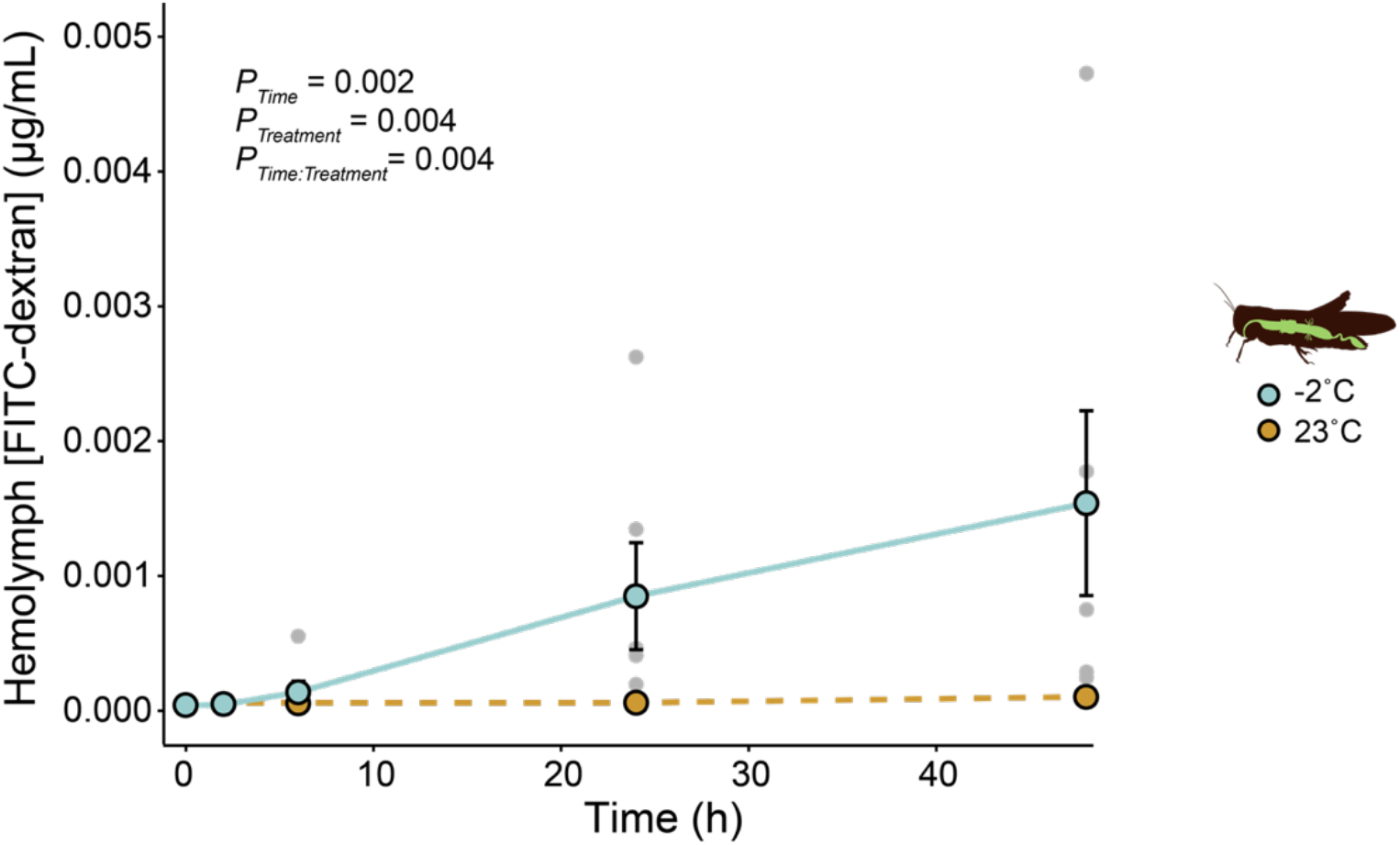
FITC-dextran leaks in the mucosal to serosal direction across the gut of *L. migratoria*. Levels of hemolymph FITC-dextran (μg/mL) following oral administration of the marker and either 2, 6, 24, or 48 h at −2°C (n = 6 per time point). The dotted line represents sampled control locusts across 48 h at 23°C (n = 6 per time point). Values are mean ± standard error. Grey points represent each sample taken per time point. Error bars not shown are obscured by the symbols.

## Discussion

Chill susceptible insects experience adverse effects of chilling at low temperatures that occur in the absence of ice formation. Consequences of cold exposure for these insects, like chill coma and tissue damage, are consistently associated with a disruption of ion and water balance (Overgaard and MacMillan, 2017). Although temperature effects on active ion transport processes are likely critical drivers of organismal failure, another potential contributor is cold-induced deterioration of paracellular barrier components known as septate junctions (SJs). In this study, we provide the first evidence for the presence of unidirectional cold-induced paracellular leak. To our knowledge, this is also the first evidence of paracellular leak in cold-exposed insects other than *Drosophila*. Our findings provide additional correlative evidence of a role for epithelial barrier function as a contributing factor in insect chill tolerance.

Similar to trends observed in *Drosophila*, we predicted that a large and rapid increase in gut FITC-dextran concentration (i.e. serosal to mucosal leak) would occur over time in the cold (MacMillan et al., 2017). However, when each gut segment was analysed for marker content, we found that cold stress induced minimal leak from the hemolymph into the gut (Fig. 2B). Even more striking was the lack of difference found between control and cold exposed locusts when the total amount of FITC-dextran within the gut at 24 h was compared (Fig. 2C). These results therefore suggest that a slight leak occurs in the serosal to mucosal direction, however, it is not temperature sensitive. Similarly, we also observed no difference between the amount of FITC-dextran in the hemolymph in the cold and at room temperature (Fig. 2D). It is important to note that while we saw decreasing levels of FITC-dextran from the hemolymph overall, the majority of the injected marker was retained in the hemocoel after a prolonged period of time. Such FITC-dextran retention within the hemolymph (whether through a lack of or minimal presence of marker degradation) has also been observed in vertebrate models like rats (*Ratticus norvegicus domestica*), as well as in invertebrate models like the plant bug (*Lygus hesperus*), lepidopterans (*Malacosoma disstria, Manduca sexta, Orgyia leucostigma*, and *Orgyia pseudotsugata*), and orthopterans (*Schistocerca americana, Melanoplus sanguinipes*, and *Phoetaliotes nebrascensis;* Barbehenn and Martin, 1995; Habibi et al., 2002; Nejdfors et al., 2000). Based on this evidence, we excluded FITC-dextran metabolism as a plausible explanation for our nominal marker movement, and remain confident that dextran is a good marker of paracellular permeability.

The lack of marker movement we observed is very small when compared to results in *Drosophila*, where even cold-acclimated (and more cold tolerant) flies exhibited 10.5-fold increases in hemolymph FITC-dextran levels in the cold (Andersen et al., 2017b; MacMillan et al., 2017). While in lesser concentrations, other macromolecules such as inulin (approx. 5000 kDa) have also been shown to leak across the gut and into the hemolymph in fifth instar desert locusts (S. *gregaria;* desert locust) (Zhu et al., 2001). Furthermore, areas along the midgut of larval *Aedes aegypti* (yellow fever mosquito) are permeable to FITC-dextran as large as 148 kDa (Edwards and Jacobs-Lorena, 2000). This permeability to large molecules is not a phenomenon limited to invertebrates. On the contrary, numerous intestinal permeability experiments have been done in vertebrate models such as mice and rats using FITC-dextran – the vast majority of which support the ability of FITC-dextran to diffuse across areas of the gut (Pantzar et al., 1993; Woting and Blaut, 2018). While our findings clearly differ from those previously reported, they are consistent with the inability for FITC-dextran to permeate the locust rectal wall (Gerber and Overgaard, 2018). In this case, however, gut preparations were exposed to a short-term cold stress. Exposure to more prolonged bouts of cold may yield different results. A plausible explanation for this lack of marker movement in our locusts may thus lie in the structure of FITC-dextran itself. Permeability across the paracellular pathway is largely determined by septate junctions (SJs), which span an intercellular space of 50-200 Å (5-20 nm; Jonusaite et al., 2016). By comparison, the 4 kDa FITC-dextran used in this study is approximately 14 Å and should therefore be able to cross through this pathway unhindered. However, as FITC-dextran is bulky, polar, and uncharged, it may be physically incapable of permeating the paracellular pathway in our locusts, even under cold stress (Matter and Balda, 2003).

While our initial experiment examined movement in the serosal to mucosal direction to isolate gut leak, it is far more common practice to assess leak via the opposite route. In addition to *Drosophila*, studies spanning an array of models from kissing bugs (*Rhodnius prolixus*) to mice (*Mus musculus*) and killifish (*Fundulus heteroclitus*) have documented the movement of various markers through the paracellular pathways in the mucosal to serosal direction across gut epithelia (Andersen et al., 2017b; le Skaer et al., 1987; MacMillan et al., 2017; O’Donnell and Maddrell, 1983; O’Donnell et al., 1984; Wood and Grosell, 2012; Woting and Blaut, 2018). The lack of FITC-dextran movement in the serosal to mucosal direction may therefore be attributed to the route of administration.

To distinguish between the presence of strictly unidirectional leak and a lack of leak, we fed locusts a dry food-dextran mixture 24 h prior to cold exposure. In stark contrast to our previous results, we observed a significant increase in hemolymph dextran concentrations over time at −2°C, resulting in a near 18- and 32-fold rise in total FITC-dextran levels after 24 h and 48 h in the cold, respectively (Fig. 3). Interestingly, the former increase of FITC-dextran under cold stress was similar to that seen across 24 h in FITC-dextran-fed *Drosophila*, where a 10.5-fold increase in marker concentration was observed (MacMillan et al., 2017). Meanwhile, in the opposite direction (serosal to mucosal), leak across the gut of our locusts resulted only in a 2-fold increase in the cold relative to concentrations in animals prior to cold exposure (Fig. 2B). These data therefore suggest that cold-induced leak does occur in locusts, but occurs unidirectionally across the gut epithelia during cold stress.

It is well-documented that cold exposure causes not only a loss of ion and water balance, but also a depolymerization of cytoskeletal components such as actin (Belous, 1992; Callaini et al., 1991; Des Marteaux et al., 2018; Kayukawa and Ishikawa, 2009; Kim et al., 2006). The actin cytoskeleton is critical to ion transport regulation, often acting as an anchor point to which transport proteins attach on both the basolateral and apical borders of epithelial cells (Cantiello, 1995a; Cantiello et al., 1991; Janmey, 1998; O’Donnell, 2017; Sasaki et al., 2014). Damage to the actin cytoskeleton can therefore create a cascade of detrimental effects within an organism. For instance, disruptions in the actin cytoskeleton have been shown to inactivate K^+^ channels in human melanoma cells (Cantiello et al., 1993). Similarly, in rat kidneys, failure of the cytoskeletal system stimulates Na^+^/K^+^-ATPase activity such that it has an increased affinity for Na^+^ (Cantiello, 1995b). Such a disruption of ionoregulation could in turn directly compromise water and Na^+^ reabsorption within insects - especially in the cold.

In addition to its role in regulating ion transport, actin has also been linked to the maintenance of tissue integrity as a key component of SJ structure (Lane and Flores, 1988; Woods and Bryant, 1991). As SJs are typically located on the apical borders of epithelial cells, cold-induced disruption in SJs, as seen with *Drosophila* (MacMillan et al., 2017), may be caused by cold-induced disassembly of the cytoskeletal network (Belous, 1992; Harvey and Zerahn, 1972). Coupled with a failure of ion transporters and channels on the apical borders, such a loss of tissue integrity could lead to a functionally “funnel-like” cavity along the mucosal side of gut epithelia. This further deterioration of barrier integrity in the cold may exacerbate the damage and leak of gut contents into the hemocoel and contribute to the cascade of events which result in the damage and death typically seen in cold-exposed insects. It is important to note that the surface of gut epithelial cells may differ in transport and SJ properties, potentially resulting in only a section or sections along the gut vulnerable to cold-induced structural damage (Cioffi, 1984; Harvey and Zerahn, 1972). Nevertheless, such structural deterioration along the mucosal side of the gut epithelia may therefore account for why FITC-dextran, despite its large and bulky composition, is able to move from the gut lumen into the hemocoel, but not in the opposite direction. It remains entirely unclear whether this unidirectional leak occurs under other conditions of stress and in other animal models.

In conclusion, locusts, like *D. melanogaster*, experience cold-induced leak from the gut when exposed to low temperatures. In locusts (and possibly other insects), this leak is unidirectional across the gut and may be attributed to a cold-induced deterioration of paracellular barrier integrity along the mucosal surface of the gut. While this study presents clear evidence supporting a directionality of cold-induced leak along the gut, the precise location of this leak and the mechanisms that drive it, unfortunately, remain unclear. To work toward filling these gaps in knowledge, we propose further investigation of the effects of cold exposure on the locust gut be conducted in isolated *ex vivo* gut sac preparations (Hanrahan et al., 1984). Such an experimental setup would allow each gut region to be more carefully assayed for the presence of cold-induced damage and/or leak. We found that there is the wide variability in the degree of paracellular barrier failure observed among individual cold-exposed locusts (Fig. 3). If barrier failure is central to the progression of chilling injury, these differences in individual leak rates may closely reflect survival outcomes (Fig 1B). Combining methods of assessing survival and FITC-dextran movement (i.e. hemolymph extractions) may therefore yield useful information regarding this individual variation, and perhaps a new method for measuring and explaining insect chill susceptibility.

## Acknowledgements

The authors wish to thank Jeffery Dawson, Jessica Forrest, and Jayne Yack for providing constructive feedback used to improve the manuscript; Jeffery Dawson for supplying locusts and the tools used to construct experimental equipment; Marshall Ritchie and Charlene Mae Herrera for taking care of the locust colony during this time.

## Competing Interests

The authors declare no competing interests.

## Funding

This work was supported by a Natural Sciences and Engineering Research Council (NSERC) Discovery Grant to H.M. (RGPIN-2018-05322).

## Data Availability

All data is provided as a supplementary file for review and the same file will be uploaded to a data repository (e.g. Dryad) should the manuscript be accepted for publication.

